# Autonomous functionality of an upstream open reading frame in polycistronic mammalian mRNAs

**DOI:** 10.1101/325571

**Authors:** Shohei Kitano, Gabriel Pratt, Keizo Takao, Yasunori Aizawa

## Abstract

Upstream open reading frames (uORFs) are established as *cis*-acting elements for eukaryotic translation of annotated ORFs (anORFs) located on the same mRNAs. Here, we identified a mammalian uORF with functions that are independent from anORF translation regulation. Bioinformatics screening using ribosome profiling data of human and mouse brains yielded 308 neurologically vital genes from which anORF and uORFs are polycistronically translated in both species. Among them, *Arhgef9* contains a uORF named SPICA, which is highly conserved among vertebrates and stably translated only in specific brain regions of mice. Disruption of SPICA translation by ATG-to-TAG substitutions did not perturb translation or function of its anORF product, collybistin. SPICA-null mice displayed abnormal maternal reproductive performance and enhanced anxiety-like behavior, characteristic of *ARHGEF9*-associated neurological disorders. This study demonstrates that mammalian uORFs can be independent genetic units, revising the prevailing dogma of the monocistronic gene in mammals, and even eukaryotes.

## INTRODUCTION

Mammalian mRNAs have on average at least one upstream open reading frame (uORF) (Calvo et al., 2009), and thousands of uORFs have strong translational activity, as recently indicated by ribosome profiling (Ribo-seq) (Ingolia et al., 2011; Ingolia et al., 2014). The central concept regarding uORF functionality has been established for almost three decades: uORFs act as *cis*-regulatory elements to modulate translation levels from annotated ORFs (anORFs) located on the same mRNA transcript (Mueller and Hinnebusch, 1986; Calvo et al., 2009; Johnstone et al., 2016). In other words, the current prevailing paradigm posits that uORFs merely assist with the functional expression of anORFs. Whether any of the translatable uORFs can elicit functions independently from anORF translation has yet to be addressed (Andrews and Rothnagel, 2014; Baboo and Cook, 2014; Couso and Patraquim, 2017).

In our search for such autonomously functional uORFs, we hypothesized that the following two criteria should be satisfied: (1) anORF translation is not affected by removal of uORF translation on the same mRNA (that is, the uORF is not a *cis*-acting element for the anORF); and (2) uORF removal induces abnormality in physiological functions. In our bioinformatics screen for uORFs that meet the former, Ribo-seq data were used to obtain mammalian mRNAs that are polycistronically translated in both human and mouse brains. Of the candidates, we focused on mRNAs from *ARHGEF9* (Cdc42 guanine nucleotide exchange factor 9). After evaluating the endogenous translational expression from one uORF (called SPICA) on *ARHGEF9* mRNAs in different mouse tissues, we then performed an *in vitro* translation reporter assay to address the second criterion: whether any mutation allowed us to remove only SPICA translation and to maintain its anORF translation at the wild-type level. The identified mutation was applied to generate a SPICA-null mouse, which was then examined to assess if the anORF translation was not affected *in vivo* by the loss of SPICA translation and then if SPICA by itself influenced any mouse phenotypes. SPICA is the first example of an autonomously functional uORF in any vertebrate. This study opens doors for our better understanding of eukaryotic genes, which have been widely recognized to be monocistronic.

## RESULTS

### Bicistronic Genes in Human and Mouse Brains

We first used Ribo-seq data for human and mouse brains (Gonzalez et al., 2014) to search for mammalian mRNAs from which uORF regions and anORFs on individual mRNAs were co-translated at equivalent levels, with the assumption that some uORFs on these mRNAs could be translated in a manner that is not coordinated with anORF translation (Figure 1A). We obtained 13,602 human and 8,246 mouse mRNAs that are translated mRNAs in the brain, to which either an anORF or merged uORF (Figure 1B), or both were mapped with reads of at least 1 read per kilobase per million mapped reads (RPKM). Of these, 4,728 human and 4,767 mouse mRNAs (2,522 human and 3,173 mouse genes, respectively) were loaded with translating ribosomes at equivalent densities on both anORFs and the merged uORFs (termed “polycistronic genes”; Figure 1, C and D).

**Figure 1.**
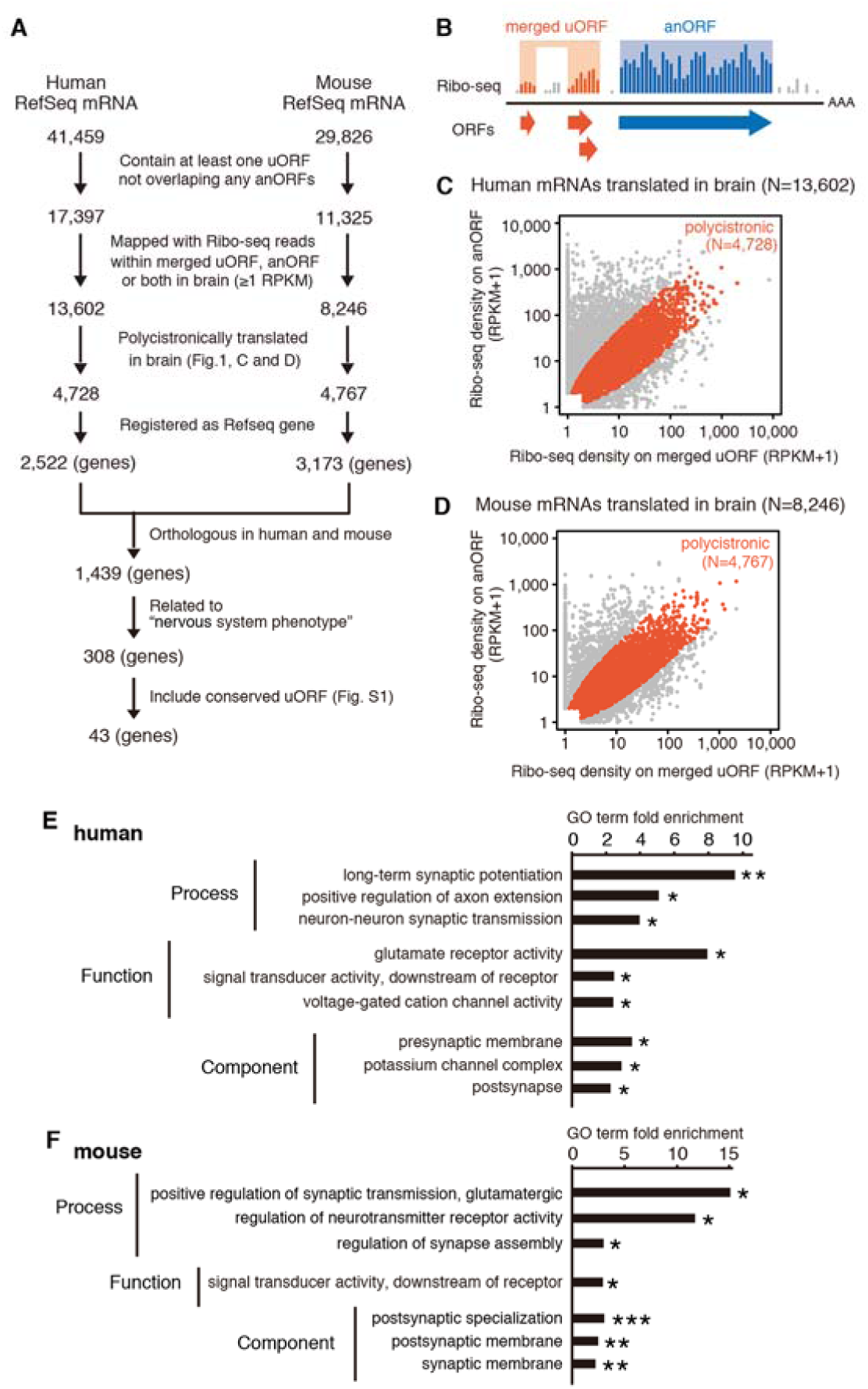
Bicistronic genes in human and mouse brains. (A) A screening scheme for candidate mRNAs with autonomously functional uORFs in the mammalian brain. (B) Definition of merged uORFs (genomic regions covered by all uORFs) and annotated ORFs (anORFs). (C and D) Scatter plots of average densities of Ribo-seq reads on merged uORFs and anORFs on individual mRNAs transcribed in (C) human and (D) mouse brains. (E and F) GO analysis of mRNAs polycistronically translated in the (E) human and (F) mouse brain. The 4,728 and 4,767 polycistronic mRNAs were compared with all 13,602 and 8,246 mRNAs translated in the human and mouse brains, respectively. \**p* < 0.001; \*\**p* < 0.0001; ***p < 1 × 10^−5^; hypergeometric test.

Gene ontology (GO) enrichment analysis showed that these polycistronic genes were significantly associated with neural GO terms in both species, especially terms related to the synapse, as compared with all the 13,602 human and 8,246 mouse mRNAs that are translated mRNAs in the brain as the background (Figure 1, E and F). This implies that some of these brain-specific polycistronic genes might be evolutionarily adapted to use uORF translation for establishing and/or regulating synaptic communication underlying brain function.

The 1,439 genes that were polycistronically translated in both human and mouse brains were then screened for genes that were previously shown to regulate brain functions by knockout mouse studies, as specified in the Mouse Genome Informatics database (Blake et al., 2009), which resulted in the identification of 308 genes. After filtering the 308 genes based on their length and conservation of amino acid sequences coded in the uORFs (≥31 codons in length and ≥70% amino acid conservation between humans and mice; Figure S1), we obtained 43 genes (Table S1).

### Region-Specific Translation of SPICA in the Brain

Among these candidates of genes encoding autonomously functional uORFs, we focused on an X-linked gene, *ARHGEF9*, that encodes collybistin in its anORF, a brain-specific GTP/GDP exchange factor (Kins et al., 2000). Collybistin participates in inhibitory synapse development by recruiting gephyrin, the main scaffolding protein in inhibitory postsynaptic densities (Grosskreutz et al., 2001; Harvey et al., 2004; Papadopoulos et al., 2007). This well-studied molecular function of collybistin has been used to explain the neurological genetic disorders associated with disruption of the *ARHGEF9* locus (Bhat et al., 2016).

One uORF of this gene, which is highly conserved among vertebrates at the encoded amino acid sequence level (Figure S2), was mapped with more Ribo-seq reads in both human and mouse brains than the anORFs (Figure 2, A and B). Our custom-made antibodies demonstrated that, like collybistin, the uORF was translated only in the brain (Figure 2C), whereas mouse *Arhgef9* mRNA was detected in the brain and heart (Figure 2D). We named this uORF “SPICA” for “<u>s</u>mall <u>p</u>rotein-<u>c</u>oding ORF in *<u>A</u>RHGEF9*”. Whereas collybistin was detected in all five of the tested brain regions at equivalent levels, the translation of SPICA was region specific, with higher protein levels in the hippocampus and cerebrum (Figure 2E). Because of a <2-fold difference in the mRNA level among these brain regions (Figure 2F), the region-specific SPICA protein levels were clearly established at the translational level. This uncorrelated translation mode of the two ORFs on the same mRNA suggested that SPICA might not be a *cis*-acting element for collybistin translation.

**Fig 2.**
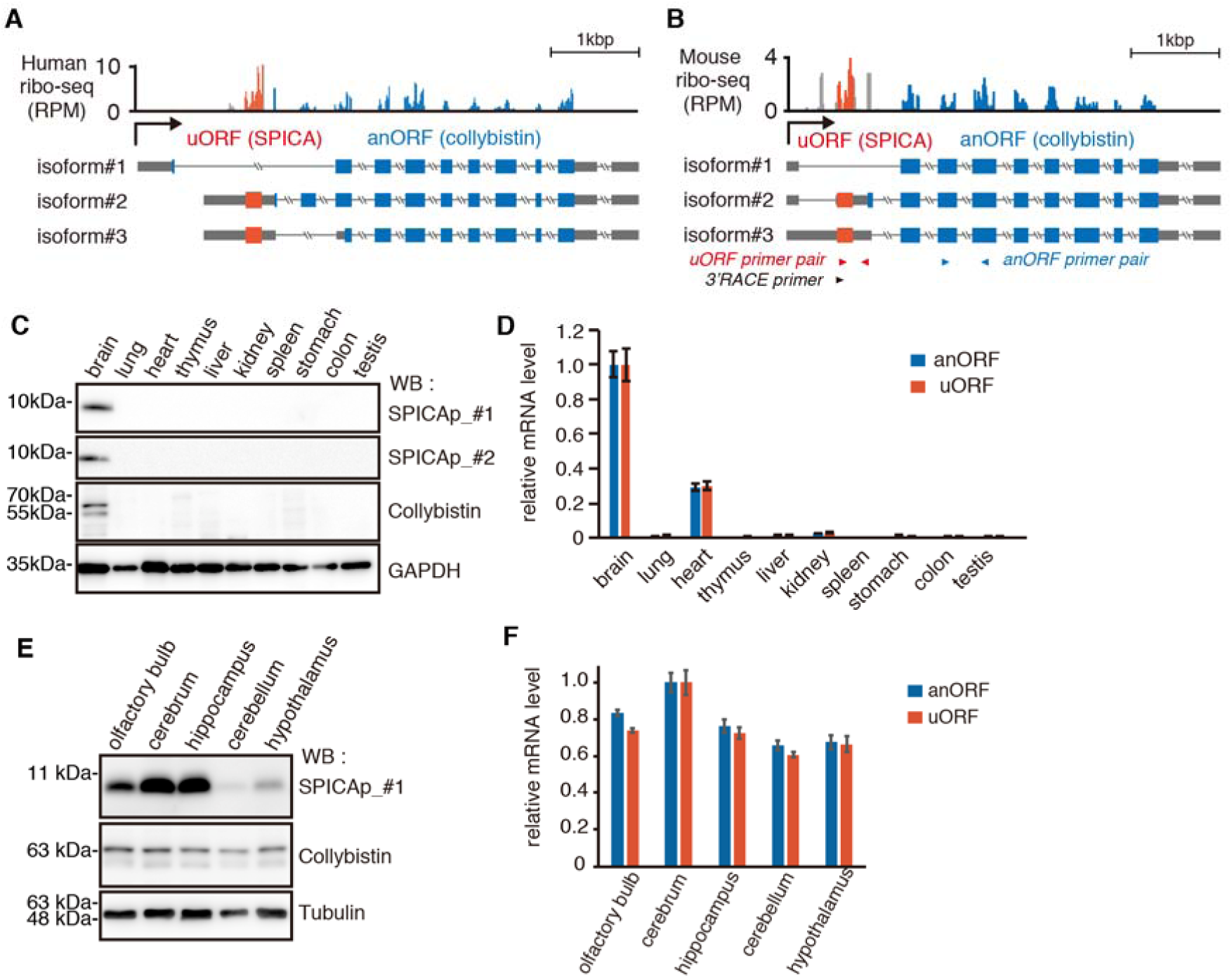
Region-specific translation of SPICA in the brain. (**A** and **B**) Ribosome occupancies on (A) human *ARHGEF9* and (B) mouse *Arhgef9* in human and mouse brains, respectively. RPM, reads per million. Primer sites for quantitative RT-PCR experiments presented in Figures 2D, 2F and 3B are indicated with arrowheads. (**C**) Immunoblots of SPICA protein and collybistin in 10 different mouse tissues. Two different anti-SPICAp antibodies were used. (**D**) Quantitative RT-PCR analysis for the anORF (collybistin) and uORF (SPICA) regions in 10 different mouse tissues. Mean ± S.E. (n = 3 biological replicates). (**E**) Immunoblots of SPICA protein and collybistin in five different mouse brain regions. (**F**) Levels of mRNA containing the collybistin ORF and SPICA in five different mouse brain regions. Mean ± S.E. (n = 3 biological replicates).

### Loss of SPICA Has Little Impact on Its anORF Translation and Functionality

To test this, we first performed *in vitro* reporter assays (Figure 3A). Wild-type (WT) plasmids of mouse *Arhgef9* mRNA isoforms #2 and #3 included their entire 5′-untranslated region (UTR) sequences, followed by a coding sequence for a FLAG–green fluorescent protein (GFP) reporter protein fusion. The FLAG-GFP coding sequence included the initial 30 bp of the collybistin ORF at the 5′**-** terminus, which allowed us to maintain the region of the mRNA that might be critical for translational initiation from its anORF. In HEK293T cells, where no endogenous *Arhgef9* mRNA is transcribed, these WT plasmids reconstituted bicistronic translation from both SPICA and the FLAG-GFP–tagged anORF (Figure 3A; the 3rd and 6th lanes from the top). Deletion of the entire SPICA region from the 5′-UTR increased FLAG-GFP translation, in agreement with the *cis*-acting model (Figure 3A; the 5th and 8th lanes). However, when we substituted the initial ATG and two internal and in-frame ATGs of SPICA with TAG stop codons, SPICA translation was expectedly lost and yet translation of the FLAG-GFP reporter was unchanged (Figure 3A; the 4th and 7th lanes from the top).

**Fig 3.**
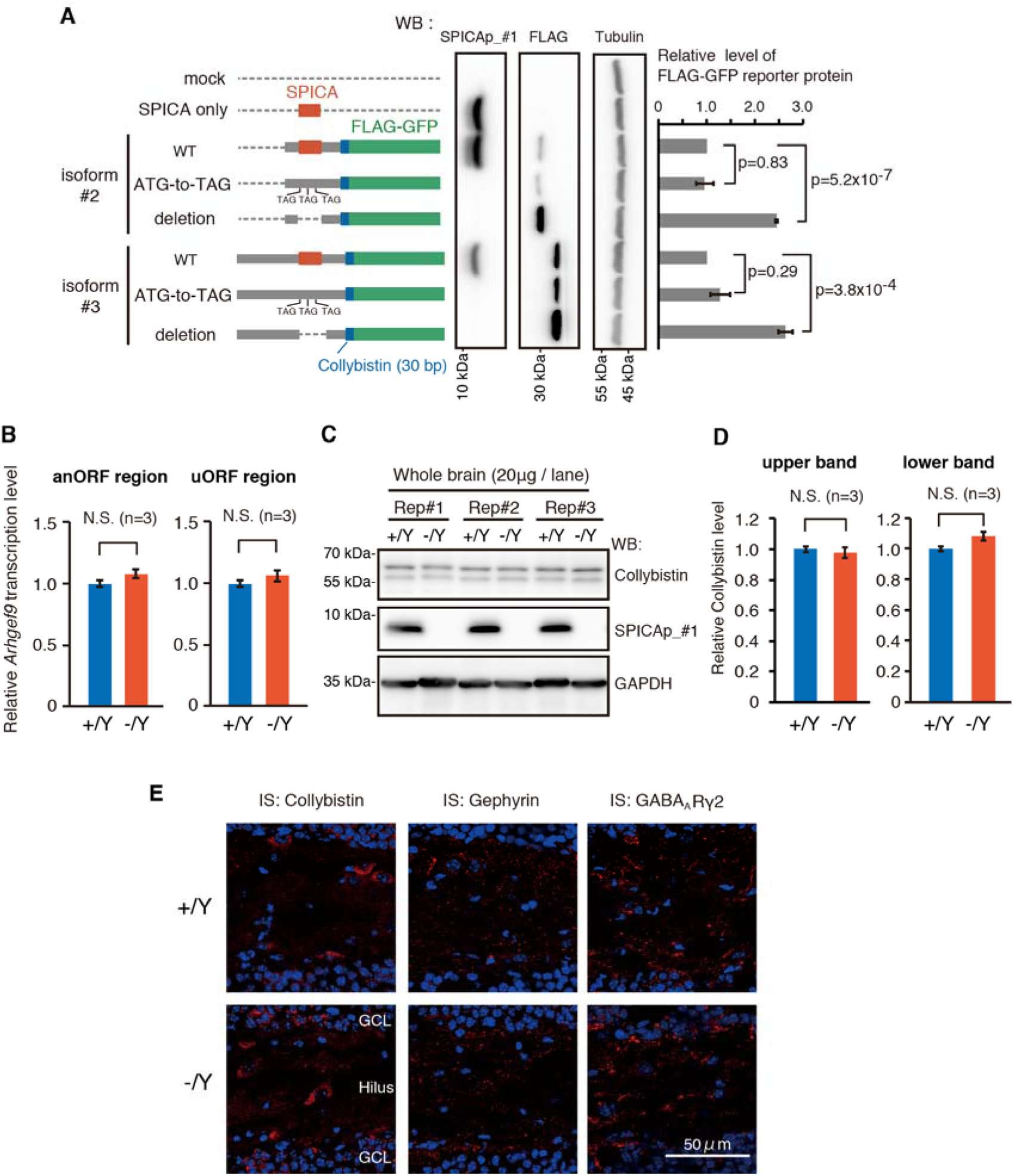
Loss of SPICA has little impact on its anORF translation and functionality. (**A**) Bicistronic translation reporter assays in HEK293T cells. The wild-type (WT) and mutant 5′-UTRs of *Arhgef9* mRNA isoforms were ligated to the FLAG-GFP fusion protein coding sequence, which starts with the first 30 bp of the collybistin anORF. Hypergeometric test. (**B** and **C**) (B) Quantitative RT-PCR and (C) immunoblots of collybistin (anORF) and SPICA (uORF) in the wild-type and SPICA-null mouse brains. Mean ± S.E. (**D**) Average intensities of the upper and lower bands of collybistin in (C) were determined using ImageJ. No significant differences in intensities were detected between wild-type and SPICA-null male littermates. Student’s t-test. (**E**) Immunostaining of collybistin, gephyrin and GABA_A_γ2 in the hippocampus hilus and granular cell layer (GCL) of WT and SPICA-null male littermates. Nuclei were stained with DAPI (blue).

Next, we applied the same ATG-to-TAG substitution mode to generate a mouse lacking only SPICA translation (Figure S3). As observed above in the *in vitro* reporter assay, male mice devoid of SPICA translation sustained *Arhgef9* transcription (Figure 3B) and collybistin translation (Figure 3C and D) at levels similar to those in the wild-type mice. Moreover, distribution of collybistin in the mutant hippocampus was maintained, and normal recruitment and distribution of gephyrin and its interacting partner, Gamma-aminobutyric-acid A receptor, gamma 2 subunit (GABA_A_γ2) (Papadopoulos et al., 2007) were observed (Figure 3E). These data support the contention that loss of SPICA translation does not abolish translation or the molecular function of collybistin *in vivo*.

### SPICA Translation Is Essential for Reproductive Behavior and Anxiety Control

This rodent model provides a unique opportunity to address the untested question: whether a uORF can have an autonomous function that is independent of anORF translation regulation. SPICA-null mice were born at the expected Mendelian ratio and appeared normal at birth (Figure 4A), albeit with significantly (∼10%) reduced body weight (Figure 4B). We also observed that homozygous mutant mothers displayed impaired nursing behaviors on day 1 after birth (Figure 4C), which led to poor survival rates of neonatal mice, in sharp contrast to the survival of pups from their WT counterparts (Figure 4D). The bimodal distribution of the survival rates in heterozygous mutants can be explained by random inactivation of one female X-chromosome, where *Arhgef9* resides. This abnormal reproductive behavior is likely to result from the loss of SPICA translation in the mothers, not in the pups, because survival rates were equivalent among pups with different SPICA genotypes (Figure 4A) that were obtained by crossing heterozygous females with hemizygous males. Additionally, SPICA translation first occurred in the brain only after postnatal day 12 (Figure 4E); therefore, loss of SPICA translation seems to have no effect in neonatal mice. The requirement of SPICA translation for reproductive behavior is consistent with the highly conserved amino acid sequence of the SPICA-encoded protein (Figure S2).

**Fig 4.**
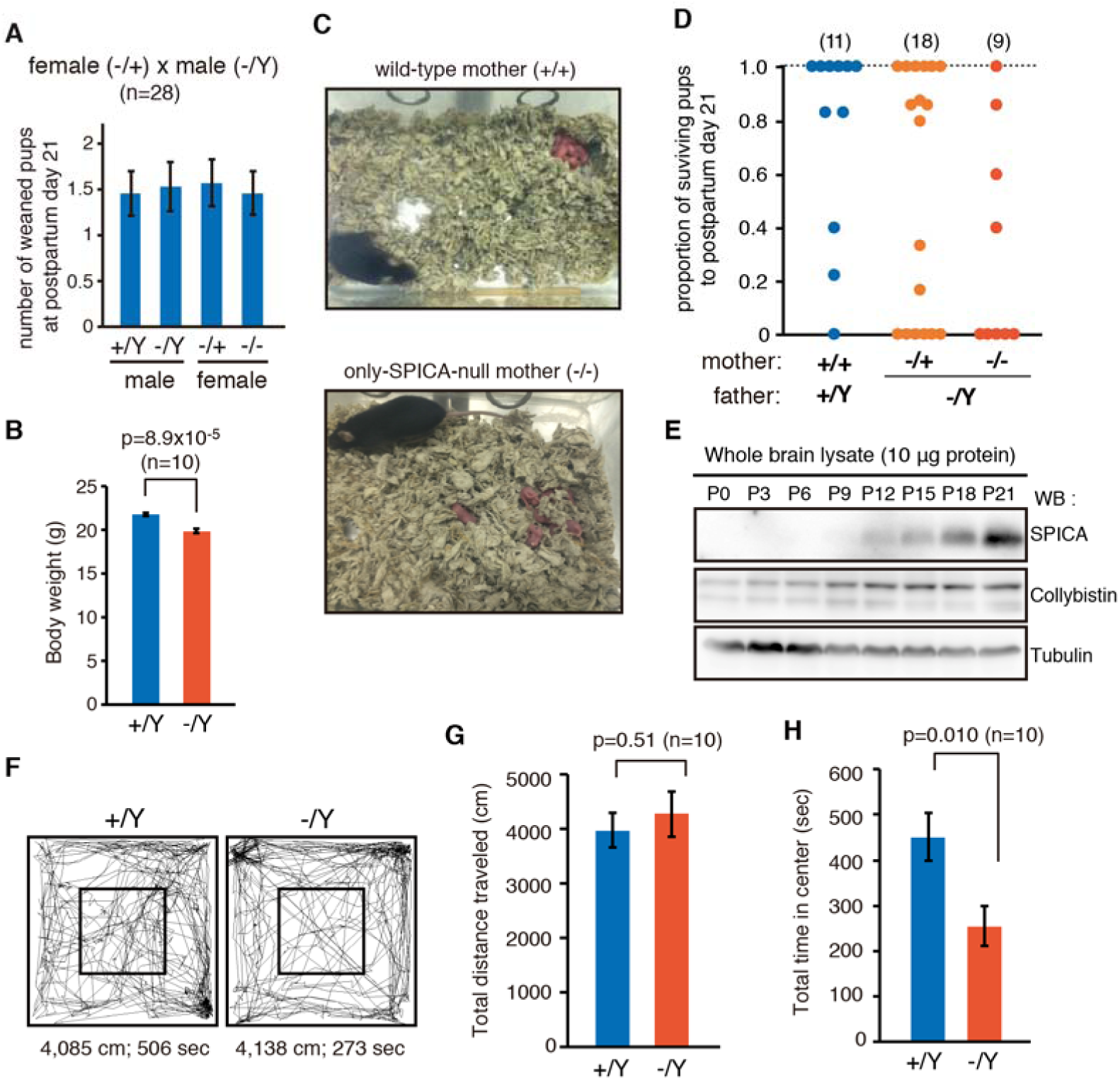
SPICA translation is essential for reproductive behavior and anxiety control. (**A**) Representative genotype data of offspring from a single heterozygous mating on postnatal day 21, showing a normal Mendelian ratio of genotypes and genders. Mean ± S.E. (**B**) Average body weights of male wild-type and SPICA-null littermates at postnatal day 56. Student’s t-test. (**C**) Typical nursing postures of wild-type and homozygous SPICA-null females on postnatal day 1. (**D**) Survival rates of pups up to postnatal day 21. The total number of matings is shown in parentheses. (**E**) Immunoblots of SPICA protein and collybistin from whole brains of neonatal mice on postnatal day (P) 0, 3, 6, 9, 12, 15, 18 and 21. **(F)** Examples of sessions from the open field test showing the walking traces of the wild-type and SPICA-null littermates. (**G** and **H**) Open field test during a 30-min period for wild-type (+/Y) and SPICA-null (–/Y) males. Student’s t-test. (G) Total distances travelled within the entire open field (40 cm × 40 cm) and (H) total time residing within the central square (18 cm × 18 cm).

To address whether this observed abnormal behavior is specific to nursing or stems from fundamental dysfunction in anxiety control, we examined the anxiety sensitivity of these mice by the open field test (Figure 4F). Previously, complete degradation of *Arhgef9* mRNA, that is, loss of both SPICA and collybistin, was shown to elevate anxiety sensitivity (Papadopoulos et al., 2007). Here, total distances walked were comparable between the wild-type and mutant male littermate pairs (Figure 4G); the latter tended to spend less time in the central area of the open field and thus had elevated levels of anxiety-like behavior (Figure 4H). These data indicate that SPICA translation by itself has a significant role in anxiety control.

## DISCUSSION

With our rationally designed experimental platform, we identified one uORF that exerts brain functions independently from regulating the translation of its own anORF. In principle, the functionality of individual ORFs on the same mRNA can be dissected only by the design of discrete mutations that eliminate only one ORF of interest without changing the translation of other ORFs on the same mRNA or affecting the mRNA transcription level. However, we speculated that this experimental approach was unlikely to be simply achieved, because it has been observed in many cases that deletion of uORFs alters translation of the downstream anORF. Therefore, our first bioinformatics screening focused on collecting uORFs that were co-translated with the anORF on the same mRNA at equivalent levels; our speculation being that translations of these uORF and anORF pairs were not correlated, as when a *cis*-acting uORF modulates anORF translation. This first screening step was performed using Ribo-seq data from human and mouse brains, which identified 1,439 genes that are bicistronically translated in the brains of both species (Figure 1A–D). This gene collection is valuable for future studies to explore the generality of autonomously functional uORFs in mammals.

This study incorporated two more gene screening steps: first, the identification of previous reports describing neural-related phenotypes after knockout of corresponding genes, and, second, the conservation among species of amino acid sequences encoded by the uORFs. The first step is useful for subsequent evaluation of phenotypes observed in engineered mice that lack only the uORF of interest. In this study, a previous report on *Arhgef9* knockout mice was informative, as it demonstrated that deletion of the fifth exon induced degradation of the entire mRNA, that is, removal of both anORF and uORF, leading to two major phenotypes: (1) impaired postsynaptic clusters of gephyrin and GABA_A_ receptors in the hippocampus and (2) abnormal anxiety control (Papadopoulos et al., 2007). Here, only the latter was observed in SPICA-null mice (Figures 3E and 4). This finding suggests that SPICA is also potentially responsible for the autism spectrum disorder noted in an individual patient that is associated with deletion of Xq11.1-11.2, which involves the SPICA-encoding exon (Bhat et al., 2016). Also, the global developmental delay diagnosed for this patient is consistent with the reduced body weight of SPICA-null mice (Figure 4B). Yet, certainly, these data do not necessarily exclude the possibility that the anORF product, collybistin, is also involved in anxiety control and normal body growth, which can be determined in mice lacking only the anORF translation in the future.

The last screening step, conservation of the amino acid sequence, was added because we expected higher sequence conservation to indicate a higher chance of detecting endogenous expression of uORF-encoded proteins. In general, conserved proteins are involved in functionally important interactions with other cellular molecules, and these interactions are likely to increase the intracellular lifespan of the interacting proteins. In our preliminary experiments, immunoblotting using anti-FLAG often failed to detect low-molecular-weight proteins tagged with FLAG (<10 kDa) even after overexpression in cell cultures. We speculated that unbound small proteins were prone to degradation in cells and, conversely, that uORF-encoded small proteins that were highly conserved and thus bound to other molecules might be expressed at a level above the detection threshold under endogenous conditions. As expected, endogenous expression of SPICA-encoded protein could be detected by immunoblotting using our custom-made antibody, enabling us to uncover the region-specific translation of SPICA in the mouse brain (Figure 2E) and to confirm the disruption of the translation of endogenous SPICA in the SPICA-null mice (Figure 3C).

These three bioinformatic steps for candidate screening were critical for the subsequent experimental demonstration of the presence of this autonomously functional uORF. To the best of our knowledge, SPICA is the first example of an autonomously functional uORF in any vertebrate. Previously, although one uORF of the C/EBPβ mRNA was deleted in mice, the authors relied on the *cis*- acting model for their experimental design and data interpretation (Wethmar et al., 2010). The SPICA/*Arhgef9* gene is a valuable model that will permit further investigation into how bicistronic mRNAs work in mammals. In particular, how two non-overlapping ORFs on one mRNA have both shared and separate roles is one of the fascinating questions to be addressed, which may help our better understanding of the structure, expression mechanism and functionality of eukaryotic genes in comparison with bacterial operons.

Like alternative splicing, autonomous functionality of uORFs is another mode representing the multi-functionality of a single gene. With this knowledge, additional caution should be taken to avoid inappropriate experimental design and/or interpretation within genetic studies. As seen for SPICA/*Arhgef9*, it is likely that deletion of only a single exon can induce degradation of the entire mRNA, resulting in removal of not only the anORF but also all the uORFs on the same mRNA. It may be worthwhile to re-examine every mutation study done in the past to determine whether uORFs, acting autonomously, may be responsible for the reported phenotypes and mechanisms.

Thanks to ribosome profiling, thousands of mammalian small open reading frames (sORFs) are now known to be translatable, and some sORF-encoded proteins have been demonstrated for long noncoding RNAs, or lncRNAs (Kikuchi et al., 2009; D’Lima et al., 2017; Huang et al., 2017; Makarewich and Olson, 2017). In contrast, for another major family of translatable sORFs, uORFs, the molecular function of any encoded proteins has yet to be demonstrated, including SPICA-encoded protein. The extremely high level of conservation of the SPICA amino acid sequence with many synonymous substitutions (Figure S2) and its expression in specific brain regions after postnatal day 12 strongly suggest the involvement of SPICA protein in neural molecular mechanisms vital for vertebrate survival. We suggest that the conceptual and experimental framework presented here will facilitate the discovery of independent functionality embedded in sORFs in the 5’-untranslated regions of mammalian and other eukaryotic mRNAs.

## EXPERIMENTAL PROCEDURES

### Bioinformatics Screening for uORFs

Human (hg19) and mouse (mm10) RefSeq mRNA BED files in the UCSC Genome Browser were used to extract all uORF sequences. The ribosome profiling data for the human and mouse brain were downloaded from NCBI SRA to quantify translation levels from the uORFs and anORFs and to screen for polycistronic mRNAs.

### RNA Purification and Quantitative RT-PCR

Total RNA was extracted from tissues from wild-type male mice using the RNeasy Mini Kit (Qiagen) and reverse-transcribed using a PrimeScript RT Reagent Kit with gDNA Eraser (TaKaRa Bio) with 1 µg of RNA. RT-PCR was performed using the Thermal Cycler Dice Real Time System (TaKaRa Bio) and SYBR Premix EX Taq II (Tli RNaseH Plus) (TaKaRa Bio). The sequences of primers used in quantitative RT-PCR are listed in Table S2.

### Antibody against SPICA-encoded protein

Anti-SPICA rabbit sera were raised against a synthetic peptide of the N-terminal region of the SPICA protein (MDSLTEQRLTSPNLPAPHLEHYSVLH). Antisera were purified with GST-fusion protein of the SPICA 5′-terminal region (corresponding to amino acid residues 2–27).

### Protein Extraction and Western Blotting

Approximately 20 μg (10 μg only for Figure 4E) of homogenates from mouse tissues was separated on a polyacrylamide gel, transferred and blotted using the following primary antibodies against SPICAp (#1: 1:1000, made in-house and #2: 1:1000, ARK Resource), collybistin (1:2000, Synaptic Systems), beta Tubulin (1:5000, Wako Pure Chemical Industries) and GAPDH (1:5000, Wako Pure Chemical Industries).

### Plasmid Construction

All the expression plasmids for cell cultures were constructed in pcDNA3 (Life Technologies) using a Gibson Assembly Master Mix (New England Biolabs).

### Cell Culture and Transfection

HEK293T cells were cultured in DMEM containing 10% FBS at 37°C under a 5% CO_2_ atmosphere and transfected with 830 ng of plasmid using 2.5 μl FuGENE-HD transfection reagent (Promega) in 12-well plates.

### Generation and Genotyping of SPICA-Null Mice

All mouse experiments were approved by the Institutional Animal Experiment Committee of the Tokyo Institute of Technology and performed in accordance with institutional and governmental guidelines. SPICA-null mice were generated by conventional homologous recombination with a floxed PGK_neo selection marker in RENKA ES cells (C57BL/6N). ES cell clones in which the PGK_neo cassette was removed by Cre were aggregated with ICR 8-cell embryos to generate chimeric mice. SPICA-null mice were backcrossed with C57BL/6NCrSlc mice three times.

### Immunostaining

Cryosections (10 μm) from the brains of postnatal day 56 male mice were incubated with primary antibody against collybistin (1:500, Synaptic Systems), Gephyrin (1:500, Synaptic Systems) and GABA_A_Rγ2 (1:1000, Synaptic Systems). The following secondary antibodies were used for detection with a confocal microscope (LSM780, Carl Zeiss): Alexa Fluor 488–conjugated goat anti–rabbit IgG or Alexa Fluor 488–conjugated goat anti–mouse IgG (1:500, Invitrogen).

### Mouse Behavioral Tests

For the open field test, male mice (postnatal day 56) were individually placed in the corner of the open field apparatus (40 cm × 40 cm × 30 cm). The total distance traveled and total time spent in the central region (18 cm × 18 cm) were recorded for 30 min per mouse. For the reproductive fitness performance test, homozygous and heterozygous SPICA-null female virgin mice were crossed with hemizygous SPICA-null males. Once pregnancy was definitive, pregnant mice were housed in separate cages from other mice. Pups that were born and survived were counted within 24 h after birth and again at postnatal day 21.

### Statistics

No statistical methods were used to predetermine sample size. For mRNA and protein analysis and mouse behavioral tests, a two-sided Student’s *t*-test was used to compare the two groups. For the reporter assay, a two-sided Student’s *t*-test was used to compare WT reporters and mutant reporters, and the significance level was corrected using the Bonferroni correction.

## ACKNOWLEDGMENTS

We thank Professor Jef D. Boeke for his valuable suggestions on this research, Mr. and Mrs. Jiro Noguchi for generous financial support for this research, Ark Resource Inc. for antibody development, J. Hirota and M. Nishida for help with mouse experiments, T. Hirano and Y. Kida for intellectual support, and the current members and alumni of the Aizawa Lab for fruitful discussions. This study was supported by grants from the Okinawa Prefectural Government (Y.A.) and JSPS KAKENHI (grant numbers 16K15116 [Y.A.] and 16J08765 [S.K.]).

## AUTHOR CONTRIBUTIONS

S.K. and Y.A. designed the experiments. S.K. conducted all the experiments with help from K.T. G.P. built the initial bioinformatic platform. S.K. and Y.A. wrote the manuscript with editing by K.T.

## Supplemental Information

Supplemental Experimental Procedures

Figures S1-S3

Tables S1 and S2

## REFERENCES

Andrews, S.J., and Rothnagel, J.A. (2014). Emerging evidence for functional peptides encoded by short open reading frames. Nat Rev Genet 15, 193–204.

Baboo, S., and Cook, P.R. (2014). “Dark matter” worlds of unstable RNA and protein. Nucleus 5, 281– 286.

Bhat, G., LaGrave, D., Millson, A., Herriges, J., Lamb, A.N., and Matalon, R. (2016). Xq11.1-11.2 deletion involving ARHGEF9 in a girl with autism spectrum disorder. Eur J Med Genet 59, 470–473.

Blake, J.A., Bult, C.J., Eppig, J.T., Kadin, J.A., Richardson, J.E., and Group, M.G.D. (2009). The Mouse Genome Database genotypes::phenotypes. Nucleic Acids Res 37, D712–719.

Calvo, S.E., Pagliarini, D.J., and Mootha, V.K. (2009). Upstream open reading frames cause widespread reduction of protein expression and are polymorphic among humans. Proc Natl Acad Sci U S A 106, 7507–7512.

Couso, J.P., and Patraquim, P. (2017). Classification and function of small open reading frames. Nat Rev Mol Cell Biol 18, 575–589.

D’Lima, N.G., Ma, J., Winkler, L., Chu, Q., Loh, K.H., Corpuz, E.O., Budnik, B.A., Lykke-Andersen, J., Saghatelian, A., Slavoff, S.A. (2017). A human microprotein that interacts with the mRNA decapping complex. Nature Chemical Biology 13, 174–180.

Gonzalez, C., Sims, J.S., Hornstein, N., Mela, A., Garcia, F., Lei, L., Gass, D.A., Amendolara, B., Bruce, J.N., Canoll, P., et al. (2014). Ribosome profiling reveals a cell-type-specific translational landscape in brain tumors. J Neurosci 34, 10924–10936.

Grosskreutz, Y., Hermann, A., Kins, S., Fuhrmann, J.C., Betz, H., and Kneussel, M. (2001). Identification of a gephyrin-binding motif in the GDP/GTP exchange factor collybistin. Biol Chem 382, 1455–1462.

Harvey, K., Duguid, I.C., Alldred, M.J., Beatty, S.E., Ward, H., Keep, N.H., Lingenfelter, S.E., Pearce, B.R., Lundgren, J., Owen, M.J., et al. (2004). The GDP-GTP exchange factor collybistin: an essential determinant of neuronal gephyrin clustering. J Neurosci 24, 5816–5826.

Huang, J.Z., Chen, M., Chen, D., Gao, X.C., Zhu, S., Huang, H., Hu, M., Zhu, H., Yan, G.R. A peptide encoded by a putative lncRNA HOXB-AS3 suppresses colon cancer growth. Mol Cell 58, 171–184.

Ingolia, N.T., Brar, G.A., Stern-Ginossar, N., Harris, M.S., Talhouarne, G.J., Jackson, S.E., Wills, M.R., and Weissman, J.S. (2014). Ribosome profiling reveals pervasive translation outside of annotated protein-coding genes. Cell Rep 8, 1365–1379.

Ingolia, N.T., Lareau, L.F., and Weissman, J.S. (2011). Ribosome profiling of mouse embryonic stem cells reveals the complexity and dynamics of mammalian proteomes. Cell 147, 789–802.

Johnstone, T.G., Bazzini, A.A., and Giraldez, A.J. (2016). Upstream ORFs are prevalent translational repressors in vertebrates. EMBO J 35, 706–723.

Kikuchi, K., Fukuda, M., Ito, T., Inoue, M., Yokoi, T., Chiku, S., Mitsuyama, T., Asai, K., Hirose, T., Aizawa, Y. (2009). Transcripts of unknown function in multiple-siganling pathways involved in human stem cell differentiation. Nucleic Acids Res 37, 4987–5000.

Kins, S., Betz, H., and Kirsch, J. (2000). Collybistin, a newly identified brain-specific GEF, induces submembrane clustering of gephyrin. Nat Neurosci 3, 22–29.

Makarewich, C.A., and Olson, E.N. (2017). Mining for micropeptides. Trends in Cell Biology 27, 685– 696.

Mishina, M., and Sakimura, K. (2007). Conditional gene targeting on the pure C57BL/6 genetic background. Neurosci Res 58, 105–112.

Mueller, P.P., and Hinnebusch, A.G. (1986). Multiple upstream AUG codons mediate translational control of GCN4. Cell 45, 201–207.

Papadopoulos, T., Korte, M., Eulenburg, V., Kubota, H., Retiounskaia, M., Harvey, R.J., Harvey, K., O’Sullivan, G.A., Laube, B., Hülsmann, S., et al. (2007). Impaired GABAergic transmission and altered hippocampal synaptic plasticity in collybistin-deficient mice. EMBO J 26, 3888–3899.

Wethmar, K., Bégay, V., Smink, J.J., Zaragoza, K., Wiesenthal, V., Dörken, B., Calkhoven, C.F., and Leutz, A. (2010). C/EBPbetaDeltauORF mice--a genetic model for uORF-mediated translational control in mammals. Genes Dev 24, 15–20.

